# A switch in kappa opioid receptor signaling from inhibitory to excitatory induced by stress in a subset of cortically-projecting dopamine neurons

**DOI:** 10.1101/2025.08.09.669424

**Authors:** Elyssa B. Margolis

## Abstract

**Background:** The kappa opioid receptor (KOR) has shown potential as a therapeutic target for several neuropsychiatric disorders including major depressive disorder, chronic pain, and substance use disorder. Signaling of G protein coupled receptors (GPCRs) like the KOR is generally thought to change in magnitude, not valence, with behavior states. Here we investigated KOR modulation of ventral tegmental area (VTA) neurons following an acute, behaviorally aversive manipulation.

**Methods:** KOR agonist responses were measured with whole cell recordings in acute brain slices containing the VTA from male and female rats. Slices were made <1 hr, 3 days or 5 days after a foot shock or sham session. Slices from untreated rats were used to determine the mechanism of action. Recordings were also made in neurons labeled by the retrograde tracer DiI to evaluate circuit specificity. Place conditioning to intra-VTA KOR agonist injections was performed in sham vs shock rats.

**Results:** After acute stress KOR activation excited a subset of VTA dopamine neurons. In slices from untreated animals, brief corticotrophin releasing factor (CRF) exposure *ex vivo* rapidly induced a similar switch in KOR signaling. This effect was observed specifically in dorsal medial prefrontal cortex (dmPFC) projecting VTA neurons, but not in other projections. Behaviorally, foot shock stress produced a loss of conditioned place aversion to intra-VTA KOR activation.

**Conclusions:** After acute aversive stress, KOR activation in the VTA excites, rather than inhibits, dmPFC-projecting VTA dopamine neurons. This change can be generated by activating the CRF system and interferes with the aversiveness of VTA KOR activation.

## Introduction

Activation of the KOR system mediates immediate processing of and long-lasting behavioral changes in response to aversive stimuli. Pharmacological blockade or genetic inactivation of KORs can decrease the acute aversiveness of stressors, including pain, and promote resilience to the development of depressive-like behaviors including decreased food reward seeking, decreased social interactions, and increased immobility in forced swim and social defeat paradigms (1–4). Therefore, KOR antagonism has been proposed as a therapeutic approach to neuropsychiatric disorders with symptom profiles that include aversive emotional and anhedonic states, such as anxiety and depression, and to mitigate stress induced relapse to drug use (5).

The opioid receptor family of GPCRs commonly signals via inhibitory Gαi/o activation (6). Agonists at somatodendritic KORs hyperpolarize neurons in a variety of brain regions including the dorsal raphe, nucleus raphe magnus, and ventral pallidum (7–9). In the VTA, KOR activation inhibits a subset of dopamine neurons through G protein-gated inwardly rectifying K^+^ channel (GIRK) activation while non-dopamine neurons show no response(10). Behaviorally, KOR activation in the VTA is aversive (11), and intra-VTA blockade of KORs prevents the aversiveness of systemic KOR agonist administration (12). Intra-VTA KOR blockade also prevents stress induced reinstatement of cocaine seeking (13). Paradoxically, aversive stress blunts some physiological KOR responses, such as the KOR coupled GIRK conductance in serotonergic dorsal raphe neurons (8). Understanding KOR function in relevant behavior states will improve our ability to treat the target neuropsychiatric disorders.

Here we investigated postsynaptic KOR agonist responses in VTA neurons after an acute aversive stressor in order to improve our understanding of what functionality KOR antagonism might have in a therapeutically relevant state. Rather than a change in response magnitude, acute aversive stress caused a change in the sign of the physiological response to KOR activation in a subset of VTA neurons. Importantly, this change was specific to VTA dopamine neurons projecting to the dmPFC, a site where dopamine function contributes to executive function and reward processing (14–17).

## Methods

### Animals

All experiments were performed in accordance with the guidelines of the National Institutes of Health Guide for the Care and Use of Laboratory Animals and the Institutional Animal Care and Use Committees (IACUC) at the University of California San Francisco. Male and female Sprague Dawley rats (p35 – p120 for electrophysiology; > 250g for behavior) were obtained from Inotiv or Charles River Laboratories. Rats were allowed access to food and water ad libitum and maintained on a reverse 12 h:12 h light/dark cycle. Rats were group housed except those with cannula implants.

### Foot shock stress

Operant chambers equipped with a foot shock grid were used for administering aversive stressors (physical boxes manufactured by Med. Associates with custom updated electronics controlled by microcontroller (Arduino)). Foot shock or sham sessions were 10 min, delivering 10, 0.8 mA intensity, 0.5 s duration shocks. Intervals between shocks were randomized (10 – 110 sec) in order to avoid predictability. Foot shock stress moderately elevated blood CORT, samples taken directly after foot shock session (Supplementary Fig. 1).

### Electrophysiology

Rats were deeply anesthetized with isoflurane, decapitated, and brains were quickly removed and cooled with ice-cold artificial cerebrospinal fluid (aCSF) consisting of (in mM): 119 NaCl, 2.5 KCl, 1.0 NaH_2_PO_4_, 26.2 NaHCO_3_, 11 glucose, 1.3 MgSO_4_, 2.5 CaCl_2_, saturated with 95% O_2_-5% CO_2_, with a measured osmolarity 310–320 mOsm/L. Horizontal sections (150 μm) containing the VTA were cut with a Leica VT 1000 S or a Campden 7000smz-2 vibratome. Tissue containing the DiI injection sites was drop fixed in 4% formaldehyde to confirm targeting, with recordings in DiI labeled neurons made blind to injection target. Slices were incubated in oxygenated aCSF at 33 °C and allowed to recover for at least one hour. A single slice was placed in the recording chamber and continuously superfused with oxygenated aCSF (2 mL/min). Neurons were visualized with an upright microscope (Zeiss AxioExaminer.D1) equipped with infrared-differential interference contrast, Dodt optics, and fluorescent illumination. Whole-cell recordings were made at 34 °C using borosilicate glass microelectrodes (3–5 MΩ) filled with K-gluconate internal solution containing (in mM): 123 K-gluconate, 10 HEPES, 8 NaCl, 0.2 EGTA, 2 MgATP, 0.3 Na_3_GTP, and 0.1% biocytin (pH 7.2 adjusted with KOH; 275 mOsm/L). Liquid junction potentials were not corrected. Input resistance was monitored throughout voltage clamp experiments with a −4 mV step every 30 s.

Signals were recorded using an Axopatch 1D (Molecular Devices, San Jose, CA) or IPA (Sutter Instruments, Novato, CA). Signals were filtered at 5 kHz and collected at 20 kHz using IGOR Pro (Wavemetrics) and custom data acquisition routines or collected at 10 kHz using SutterPatch software (Sutter Instruments).

Baseline hyperpolarization-activated cation currents (*I*_h_) were recorded in each neuron by voltage clamping cells at −60 and stepping to −40, −50, −70, −80, −90, −100, −110, and −120 mV. Recordings were then made in current clamp mode (I = 0 pA). Changes in *I*_h_ properties were measured in voltage clamp, alternating between two step series (Supplementary Fig. 2) with one series commencing each 2.17 min.

After recordings, slices were drop fixed in 4% PFA for at least 2 h at 4 °C and processed for tyrosine hydroxylase (TH) immunocytochemistry and biocytin labeling. Recording were made from throughout the VTA.

### Immunocytochemistry

Slices were preblocked for 2 h at room temperature in PBS with 0.3% Tween20 and 5% normal goat serum, then incubated at 4°C with a rabbit anti-TH polyclonal antibody (1:400; EMD Millipore, RRID: AB_390204). Slices were then washed thoroughly in PBS with 0.3% Tween20 before being agitated overnight at 4°C with Cy5 anti-rabbit secondary antibody (1:1000; Jackson ImmunoResearch, RRID: AB_2534032) and FITC streptavidin (1:200; Jackson ImmunoResearch, AB_2337236). Tween20 was excluded in slices with DiI. Sections were rinsed and mounted on slides using VECTASHIELD^®^Antifade Mounting Media (Vector Laboratories) and visualized with an Axioskop FS2 Plus (Zeiss) with an Axiocam MRm (Zeiss) running Neurolucida (MBF Biosciences).

### CORT ELISA

Tail vein blood samples were collected immediately after foot shock or sham sessions from behavior animals. Trunk blood was collected at time of slice preparation for electrophysiology animals. Samples were centrifuged, then serum was frozen and stored at −80°C. ELISA kits were used as directed (DetectX Corticosterone Enzyme Immunoassay Kit (product # K014-H1); Arbor Assays).

### Stereotaxic surgeries

#### Tracer injections

Rats weighing 275–300 g were anesthetized with 3–5% isoflurane (Henry Schein) via inhalation and secured in a stereotaxic frame. Bilateral craniotomies were created with a dental drill above the injection site.

#### Tracer injections

Bilateral injections of DiI (7% in ethanol; Biotium, Hayward, CA) were made in the dmPFC (2.0 mm AP, ± 0.6 mm ML, −2.8 mm V), vmPFC (2.0 mm AP, ± 0.6 mm ML, −4.8 mm V), NAc (1.2 mm AP, ± 0.8 mm ML, −7.4 mm V), or basolateral amygdala (−3.3 mm AP, ± 5.0 mm ML, −8.4 mm V) using a Nanoject II (Drummond Scientific, Broomall, PA). 50 – 300 nL was injected in each hemisphere over ∼5 min. The injector tip was left in place for at least 2 minutes after injection to decrease backflow and spread to tissue dorsal to the injection site. Recordings were made 7-8 days later.

#### Cannulation

For microinjections into the VTA, bilateral guide cannulae were implanted 1 mm above the VTA (−5.8 mm AP, ±0.5 mm ML,−7.2 mm DV). A dummy stylet was inserted to maintain patency of the cannulae. Cannulae were anchored with flat point screws and dental cement.

#### Analgesia and recovery

Rats were treated with subcutaneous Carprofen (5 mg/kg, Zoetis) and topical 2% lidocaine (Phoenix Pharmaceutical, Inc.) during the surgery for pain control. After surgery, animals had access to liquid Tylenol (∼1:40) in their drinking water for 3–5 days or were administered Meloxicam (s.c. 2 mg/kg, Pivetal) once per day for two days. All DiI and cannulae placements were histologically verified postmortem.

#### Conditioned place preference

Rats were allowed to recover for 1–2 weeks prior to behavioral testing. First, all rats were tested for bias in the conditioning boxes. The conditioning boxes (physical boxes purchased from Med. Associates, Georgia, VT, USA with custom updated electronics (Arduino; IR beam breaks (Adafruit)) and software have three divisions: two conditioning chambers (25 cm × 21 cm × 21 cm) with distinct visual (horizontal vs. vertical stripes) and textural (thick vs. thin mesh flooring) cues, separated by a smaller gray neutral chamber (12 cm × 21 cm × 21 cm). Animals were allowed up to three opportunities to show neutrality across the chambers during 20-min baseline sessions. Rats that displayed a consistent baseline preference (>65% of time spent in one chamber) were excluded from the study. Rats were pseudorandomly assigned to receive U69,593 in one of the conditioning chambers, and assignments were counter-balanced for each cohort.

The paradigm commenced with a foot shock or sham treatment as described above, completed in a different room from the conditioning chambers. Starting the next day, conditioning pairings (20 min) occurred once daily for 4 consecutive days alternating U69,593 (0.2 mM) and vehicle (0.5 mL/side). This 5 day series (including foot shock/sham) was repeated a second time for a total of 4 pairings for U69,593 and vehicle, each. Finally, on test day, rats were drug free and had access to all three chambers for 20 min. Difference score was defined as (Time spent in U69,593 paired chamber)−(Time spent in vehicle paired chamber).

#### Microinjections

Rats were lightly restrained in a cloth wrap for intracranial microinjections. A bilateral 33 G microinjector (PlasticsOne) that extended 1 mm ventrally beyond the guide cannula was inserted to deliver drug. Hamilton syringes were driven by a dual syringe pump to deliver the injection volume over 2 min. Injectors were left in place for 1 min after injection.

#### Brain removal and histochemistry

To check cannulae placements, rats were deeply anesthetized with an intraperitoneal injection of Euthasol (0.1 mg/kg, Virbac Animal Health) after 0.3 µL of Chicago Sky Blue (in 2% PBS) was injected through the cannulae to mark injection locations. After becoming unresponsive to noxious stimuli, the rats were transcardially perfused with 400 mL of saline, followed by 400 mL of 4% paraformaldehyde (PFA) in 0.1 M phosphate buffer. The brains were extracted and immersion-fixed in PFA for 2 h at room temperature (RT), washed two times with PBS to remove excess PFA, and stored in 1X PBS at 4°C.

Perfused brains and drop fixed DiI containing tissue were sectioned (50 μm) using a vibratome (Leica VT 1000S). Alternating slices were Nissl stained. Sections were mounted on slides using VECTASHIELD^®^ mounting medium (Vector Laboratories). Microinjection locations for rats included in the behavior study are summarized in Supplementary Fig. 3.

### Data analysis

The magnitude of U69,593 induced changes in V_m_ were quantified by comparing the mean values over the last 4 min of U69,593 application to the 4 minutes of baseline just prior to starting the drug application. Responses were tested for significance by statistically comparing 30 sec bins during these two time periods (unpaired Student’s t-test). *I*_h_ magnitude was measured as the difference between the initial capacitive response to a voltage step from −60 to −120 mV and the final current during the same step. Reported baseline Ri and firing rate are the average measurements over the first 2 min of *I*_clamp_ recording. For AP duration measurements, at least 8 waveforms were averaged together and duration was calculated from threshold on the ascending phase to recrossing threshold on the falling phase. Threshold was identified as the first datapoint where the derivative of the waveform exceeds 10 V/s. For evaluation of KOR agonist induced changes in *I*_h_ properties, see measurements described in Supplementary Fig. 2. Population statistics were performed in R. Where datasets failed tests of normality or homogeneity of variance, non-parametric permutation analyses (https://github.com/eb-margolis-neuroscience-lab/R-general-Margolislab/tree/main) were used to test for statistical significance. For behavior data, mixed measures two-way ANOVA was used with Bonferroni correction for multiple comparisons.

## Results

We performed *ex vivo* whole cell recordings in rat VTA neurons after a single session of aversive foot shock stress, making slices 30-60 min after the stressor. All statistics are described in Table 1. In addition to VTA neurons that responded to the KOR selective agonist U69,593 (1 μM) with a hyperpolarization or no change in membrane potential similar to prior observations in untreated animals (10,18,19), we unexpectedly observed depolarizations in a distinct subset of neurons from both male and female rats (Fig. 1A-D). This includes neurons confirmed as dopaminergic with immunocytochemistry (Fig. 1B, Supplementary Fig. 4A; recording locations in Supplementary Fig. 4B). Depolarizations were still observed 3 (4/18 neurons), but not 5 (0/7 neurons), days after the foot shock stress session (Fig 1d). Action potential (AP) threshold was more depolarized in neurons from foot shock stressed rats compared to shams, but the stressor did not alter other basal properties of VTA neurons including baseline input resistance (Ri), AP duration, AP peak, h current magnitude, or spontaneous firing rate (Supplementary Fig. 5). In the recordings from stressed animals, initial Ri was greater in neurons that subsequently showed a depolarization in response to U69,593 compared to neurons that subsequently showed a hyperpolarization (Supplementary Fig. 6). No other differences were detected in baseline physiological properties across VTA neurons grouped by their U69,593 responses (Supplementary Fig. 6). One possibility is that the aversive stressor decreases GIRK function in a subset of neurons, unmasking the observed depolarizing effect of U69,593. However, when we blocked GIRKs in slices from control, untreated animals with BaCl_2_ (100 μM), the distribution of responses to U69,593 was clearly different from those from stressed animals with no depolarization > 0.8 mV (Fig. 1D,E). Dopamine D2 receptor activation hyperpolarizes a subset of VTA neurons through GIRK channel activation (20). If acute, aversive stress decreases GIRK function in VTA neurons rather than change KOR signaling, we would expect to observe a shift in responses to the D2R agonist quinpirole (1 μM) as well. The means and distributions of quinpirole responses were not different between VTA neurons from sham and foot shocked rats (Fig. 1F).

**Figure 1.**
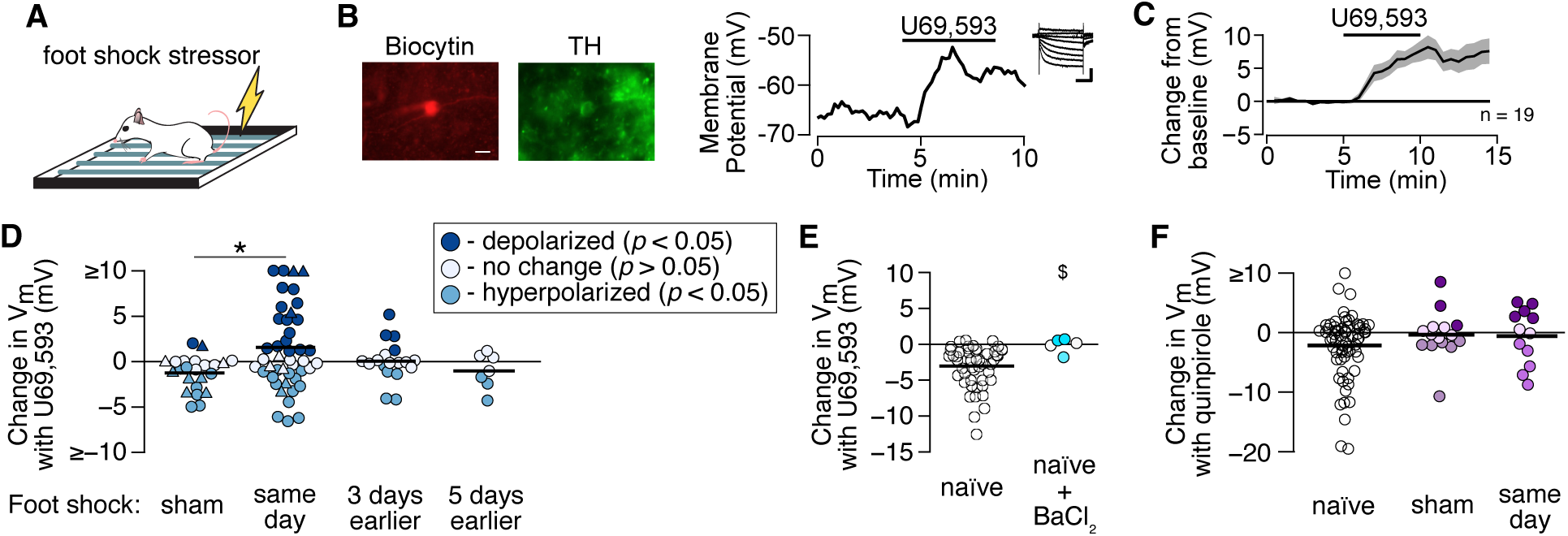
Following a single foot shock session, a subset of VTA neurons depolarize in response to KOR activation. (A) Rats experienced a single intermittent foot shock session (10 min) prior to ex vivo whole cell recordings. (B) Example recording from a VTA neuron (biocytin fill: red) that was confirmed as dopaminergic with TH immunocytochemistry (green), scale bar 10 μm. In whole cell current clamp configuration (I = 0 pA) bath application of the KOR agonist U69,592 (1 μM) caused a depolarization. Inset: I_h_ measured in voltage clamp, scale bars 200 pA and 200 ms. (C) Mean time course of responses in neurons that were significantly depolarized by U69,593. (D) Summarized data, each circle is a neuron, from same day sham, same day shock, 3 days after shock, and 5 days after shock treated rats. Circles and triangles indicate VTA neurons from male and female rats, respectively. Permutation analysis comparing means of U69,593 responses in sham vs shock, p = 0.02 (Table 1) (E) Summarized U69,593 responses in VTA neurons from naïve rats, and from naïve rats where U69,593 responses were measured in BaCl_2_ (100 μM). Permutation analysis comparing variances (SDs; $ indicates p<0.05), p = 0.04. (F) Summarized responses to the dopamine D2 receptor agonist quinpirole (1 μM) in VTA neurons from naïve, sham, and same day foot shocked rats. One way ANOVA sham vs shock, p = 0.9.

**Table 1.**
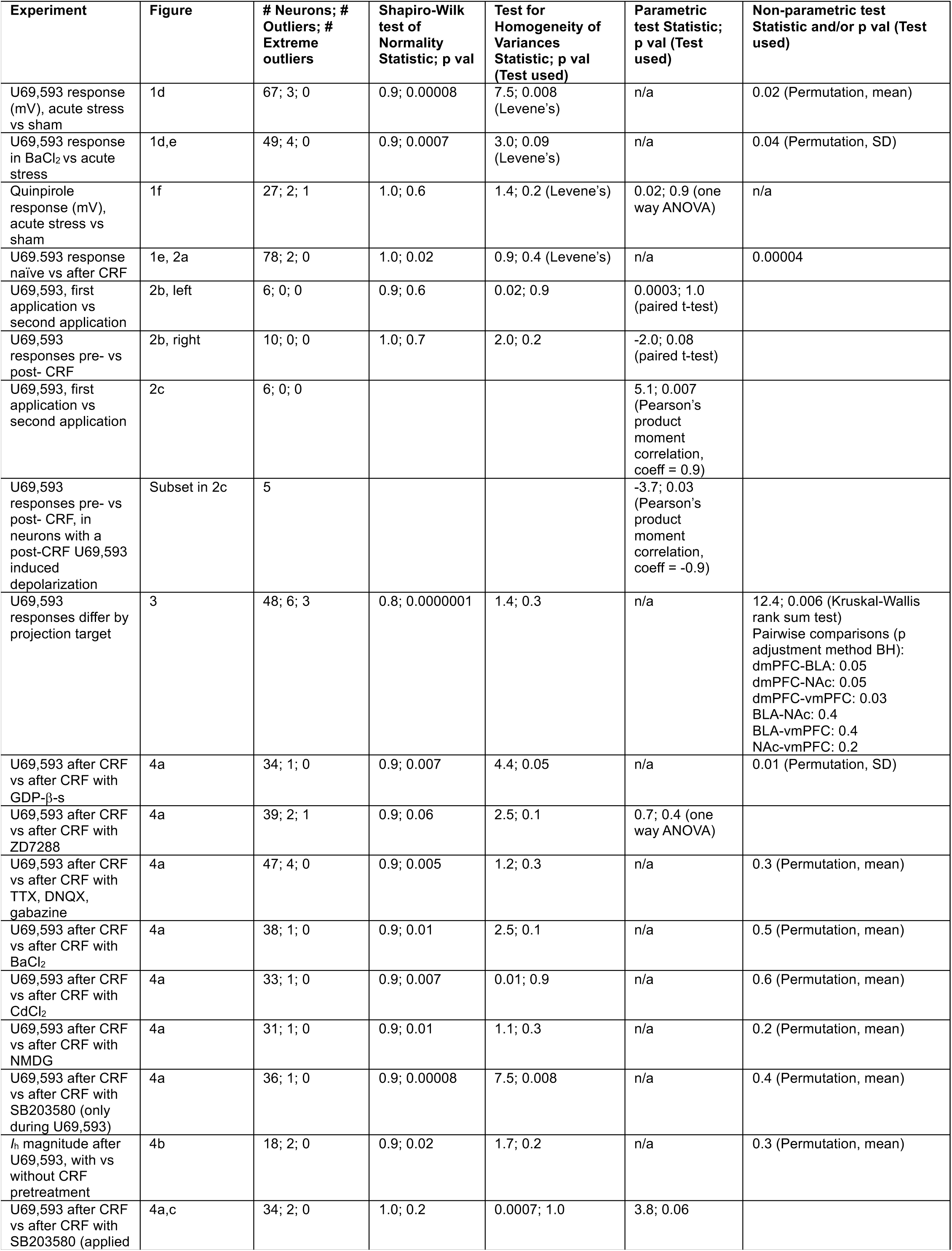
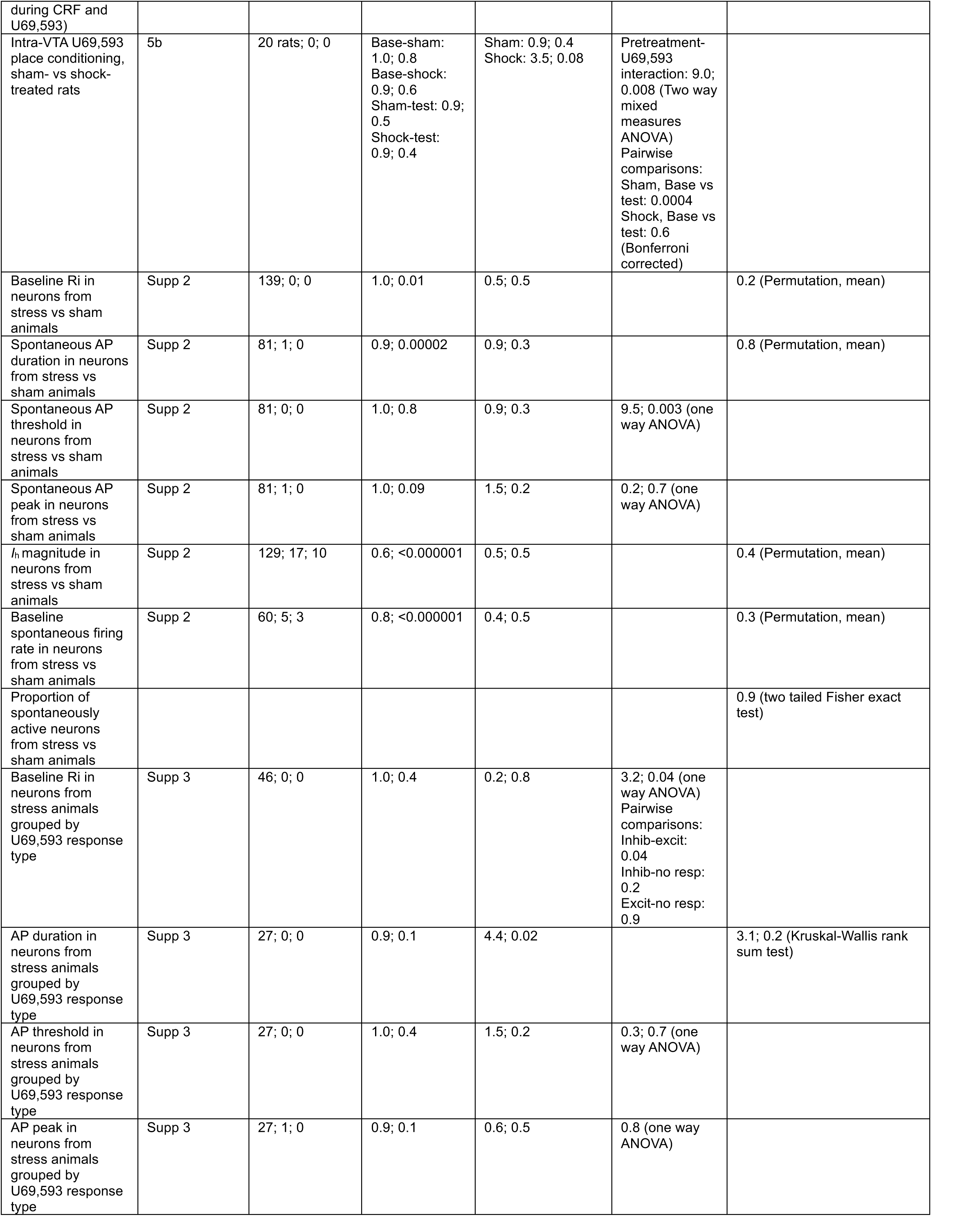
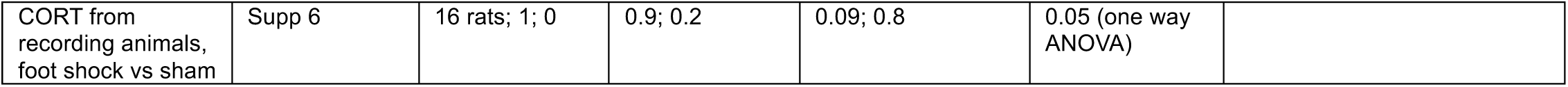
Assumption testing and statistical comparisons for summary data.

The corticotrophin releasing factor (CRF) system is upstream of the KOR system in encoding the aversiveness of stressful experiences (21), and blocking KORs can prevent CRF-induced anxiety-like behaviors (22,23). As foot shock stress drives CRF release in the VTA (24), we tested the hypothesis that incubating VTA slices from control animals with no behavioral manipulations in CRF would produce similar changes in KOR signaling to the *in vivo* aversive stressor exposure. In fact, in VTA slices incubated in 200 nM CRF for 5 min, a subset of neurons responded to subsequent U69,593 application with a depolarization (Fig. 2A). Next we tested whether neurons excited by KOR activation after CRF were the same neurons inhibited by KOR activation under control conditions or a previously KOR-insensitive population. All neurons that showed a depolarization in response to KOR activation after CRF showed a hyperpolarization to KOR activation before CRF (Fig. 2B,C). In fact, for neurons that showed a depolarization after CRF, there was a clear inverse correlation (n = 5 neurons; coeff. = −0.9) between the magnitudes of KOR responses before and after CRF application (Fig. 2C).

**Figure 2.**
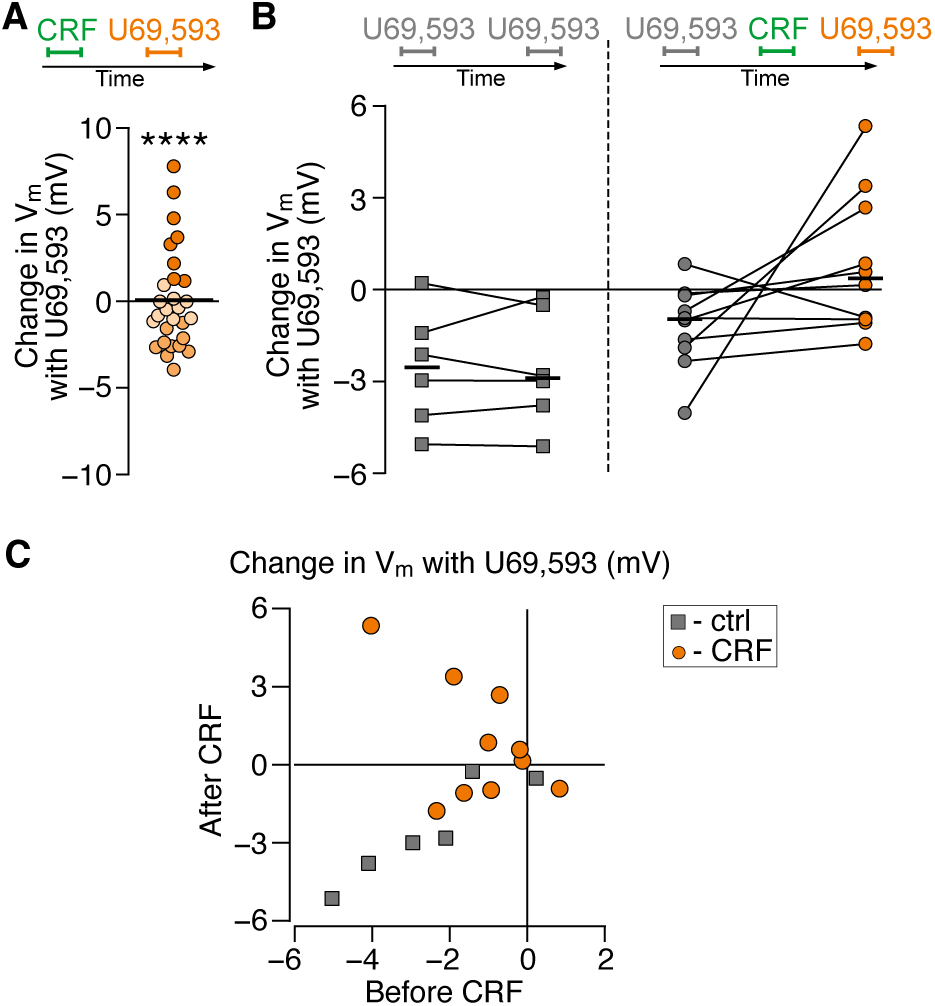
Ex vivo exposure to CRF can switch KOR signaling from inhibitory to excitatory. (A) Summarized data, each circle is a neuron. Slices were incubated for 5 min with CRF (200 nM), CRF was washed out for at least 5 min, then U69,593 (1 μM) was bath applied for 5 min. Statistical comparison to U69,593 responses in VTA neurons from naïve rats (Fig. 1E left), Permutation analysis of means, p = 0.00004. (B) Left, U69,593 (1 μM) was applied once, washed out, then reapplied (paired t-test p = 1.0). Right, U69,593 (1 μM) was applied once, washed out, CRF was washed on (5 min, 200 nM), washed out, then U69,593 was reapplied (paired t-test p = 0.08). (C) Data from (B) graphed as magnitude of response to first U69,593 application to second application.

We previously reported that VTA dopamine neurons that project to the mPFC and basolateral amygdala (BLA), but not the nucleus accumbens (NAc), are inhibited by somatodendritic KOR activation in rat (18,19). To test whether the signaling switch happens in a projection selective manner, we recorded from retrogradely labeled VTA neurons that project to dmPFC (including prelimbic and anterior cingulate cortices), ventral mPFC (infralimbic cortex), medial NAc, and BLA blind to injection site. Most distinctly, dmPFC-projecting VTA neurons showed the strongest depolarizations after CRF exposure while none of the vmPFC-projecting neurons depolarized (Fig. 3). NAc- and BLA-projecting showed no and minimal depolarizations, respectively (Fig. 3). Thus, the switch in KOR signaling from inhibitory to excitatory occurs in a specific subpopulation of dmPFC-projecting VTA neurons.

**Figure 3.**
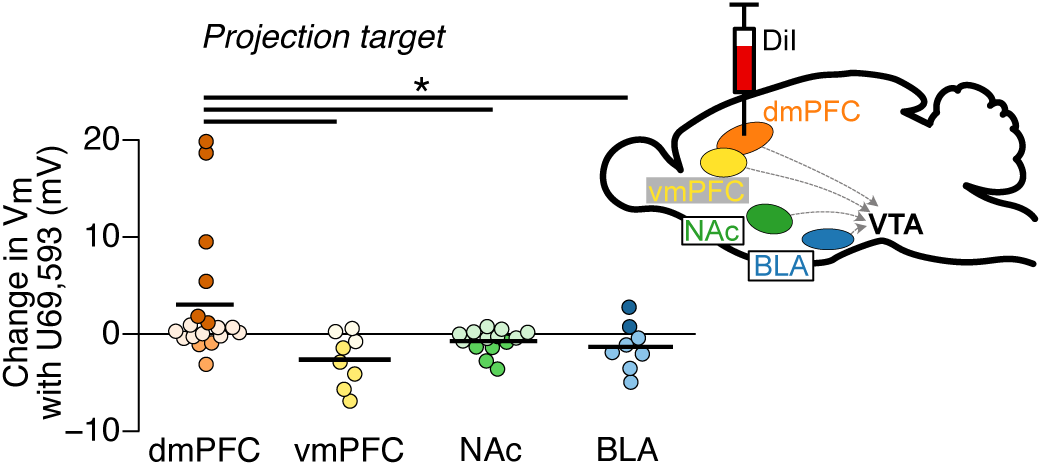
CRF switches KOR signaling in a subpopulation of dmPFC-projecting VTA neurons. U69,593 responses were recorded in retrogradely labeled neurons after slices were incubated in CRF (5 min, 200 nM). Kruskal-Wallis rank sum test p = 0.006, pairwise comparisons listed in Table 1.

Next we probed the signaling mechanism underlying the KOR mediated excitations triggered by CRF exposure. To test whether the KOR excitations are due to indirect effects involving local connectivity in the slice we measured KOR agonist responses while blocking AMPARs (10 μM DNQX), GABA_A_Rs (20 μM gabazine), and action potential activity (500 nM TTX). This did not prevent U69,593 induced depolarizations (Fig. 4A). To block G protein signaling in the recorded neuron we included GDP-β-s in the intracellular solution (500 μM), and broke in to whole cell configuration only after CRF application was completed. This G protein block eliminated both depolarizations and hyperpolarizations induced by U69,593 (Fig. 4A). Together these data show that the KOR induced depolarizations are not local circuit dependent, and because the G protein blockade was limited to the recorded neuron the KORs in question appear to be expressed in the recorded neuron. GPCR activation could cause depolarization by closing GIRKs (25), however blocking GIRKs with BaCl_2_ (100 μM) did not prevent U69,593 induced depolarizations (Fig. 4A). We previously reported that mu and delta opioid receptors (MORs and DORs) on VTA neurons can drive depolarizations that require Ca^2+^ channel conductances (26,27), however in the present study, blocking Ca^2+^ channels with bath application of CdCl_2_ (100 μM) did not prevent U69,593 induced depolarizations (Fig. 4A). A Na^+^ leak current contributes to the maintenance of a depolarized potential and spontaneous AP firing in VTA neurons (28),(29) that can be inhibited by Gαi/o protein coupled receptors (30). Blocking this conductance did not prevent U69,593 induced depolarizations (Fig. 4A).

**Figure 4.**
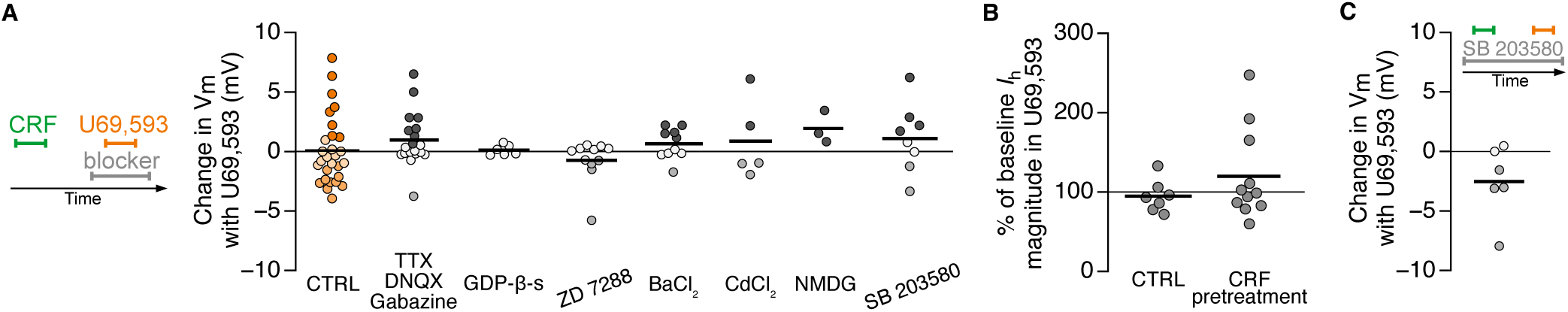
KOR mediated depolarizations require I_h_. (A) Signaling mechanism(s) that contribute to U69,593 (1 μM) induced depolarizations were investigated after exposing slices ex vivo to CRF (5 min, 200 nM). TTX (500 nM), DNQX (10 μM), and gabazine (20 μM) blocked synaptic transmission; intracellular GDP-β-s (500 μM) blocked intracellular G protein signaling; ZD7288 (10 μM) blocked HCN channels (I_h_); BaCl_2_ (100 μM) blocked K^+^ channels; CdCl_2_ (100 μM) blocked Ca^2+^ channels; replacement of Na^+^ with NMDG (151 mM, replacing NaCl) in aCSF blocked Na^+^ leak current; SB203580 (10 μM) blocked p38/MAPK signaling. (B) I_h_ magnitude was measured during U69,593 (1 μM) application in voltage clamp, stepping from −60 mV to - 120 mV, in control slices or slices pretreated with CRF (5 min, 200 nM). (C) p38/MAPK was blocked with SB203580 application during both CRF and U69,593 application.

*I*_h_ is a hyperpolarization activated, non-selective cation conductance that we found is prominent in mPFC-projecting VTA neurons (19). In nodose ganglion neurons MOR activation inhibits *I*_h_ (31), and in the nucleus raphe magnus (NRM) KOR activation augments *I*_h_ (32). The *I*_h_ blocker ZD7288 prevented U69,593 induced depolarizations (Fig. 4A). Given that the reversal potential for *I*_h_ is depolarized compared to typical membrane potentials, the simplest possibility is that KOR activation augments *I*_h_ conductance. Consistent with this, we found that U69,593 increased *I*_h_ amplitude in a subset of neurons recorded in slices pretreated with CRF, and this response that was not observed in control slices (Fig. 4B). CRF pretreatment did not impact other *I*_h_ properties measured during U69,593 application (Supplementary Fig. 2).

Bruchas and colleagues reported that P38 MAPK contributes to some KOR mediated behavioral changes following aversive stressors *in vivo* (33). Here we tested for a contribution of p38 to this KOR switch or KOR signaling. Incubating slices from control rats with the p38 blocker SB203580 prior to CRF application prevented subsequent U69,593 induced depolarizations (Fig. 4C). However, when slices were exposed to CRF prior to incubation in SB203580, U69,593 induced depolarizations persisted (Fig. 4A). We conclude that CRF receptor activation alters the signaling of a subset of KORs in the VTA through a p38 dependent signaling pathway, but KORs are not signaling through p38 directly to depolarize neurons.

Finally, we tested for a behavioral consequence of this change in KOR signaling. Decreases in dopamine neuron activity canonically encode punishment and aversion, and therefore intra-VTA KOR agonist induced aversion is widely thought to be due to inhibition of dopamine neurons. In the context of this model, a switch in KOR signaling in dopamine neurons from inhibitory to excitatory would be predicted to diminish conditioned place aversion (CPA) to intra-VTA KOR agonist microinjection. Since we found the switch in signaling sign to last between 3 and 5 days, we designed a paradigm where the animals received foot shock on days 1 and 6 in order to maintain stress-induced changes to KOR signaling. Pairings of either intra-VTA U69,593 (0.2 mM, 0.5 mL/side) or vehicle once per day were performed on non-stress days, for a total of four pairings per side (Fig. 5A), in order to minimize the formation of a direct association between the foot shock stress and the place preference manipulation. Indeed, foot shock stress pretreatment eliminated the development of CPA to intra-VTA U69,593 (Fig. 5B, placements in Supplementary Fig. 3).

**Figure 5.**
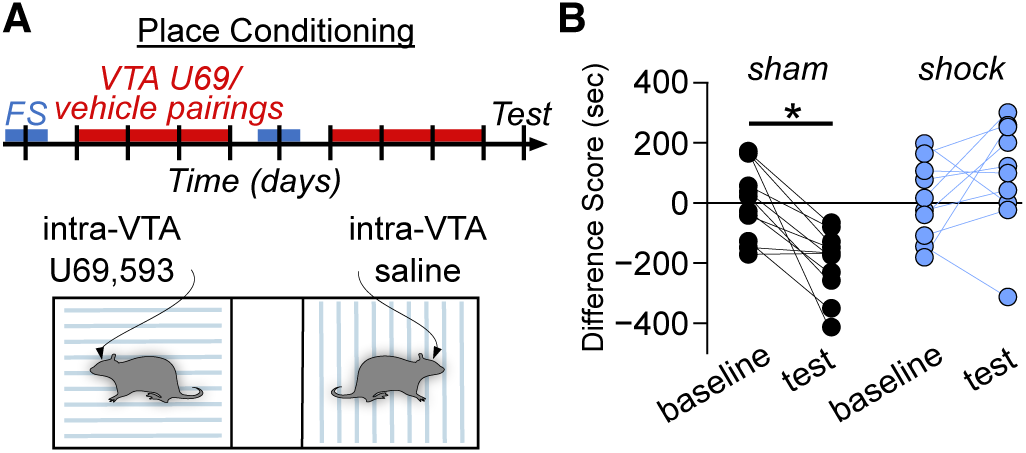
Foot shock stress eliminates intra-VTA KOR mediated aversion. (A) Experiment timeline where rats were trained in a conditioned place preference paradigm, to intra-VTA U69,593 (0.2 mM, 0.5 mL/side) or vehicle. Rats were pretreated with sham handling or foot shock stress on non-pairing days. (B) Baseline vs post training test for place conditioning. Two way mixed measures ANOVA) Pairwise comparisons: Sham, baseline vs post test: 0.0004 Shock, baseline vs post test: 0.6 (Bonferroni corrected).

## Discussion

Together these data show that a single stressful event can dramatically change the signaling sign of a modulatory GPCR system. The change occurs in a specific neural circuit and can persist for days. This stress induced switch at KORs can be induced by the stress-associated CRF neuropeptide-GPCR system in the same brain region, and while the signaling is still G protein mediated, it appears that the key conductance involved is *I*_h_, a switch from GIRK coupling (Fig. 6).

**Figure 6.**
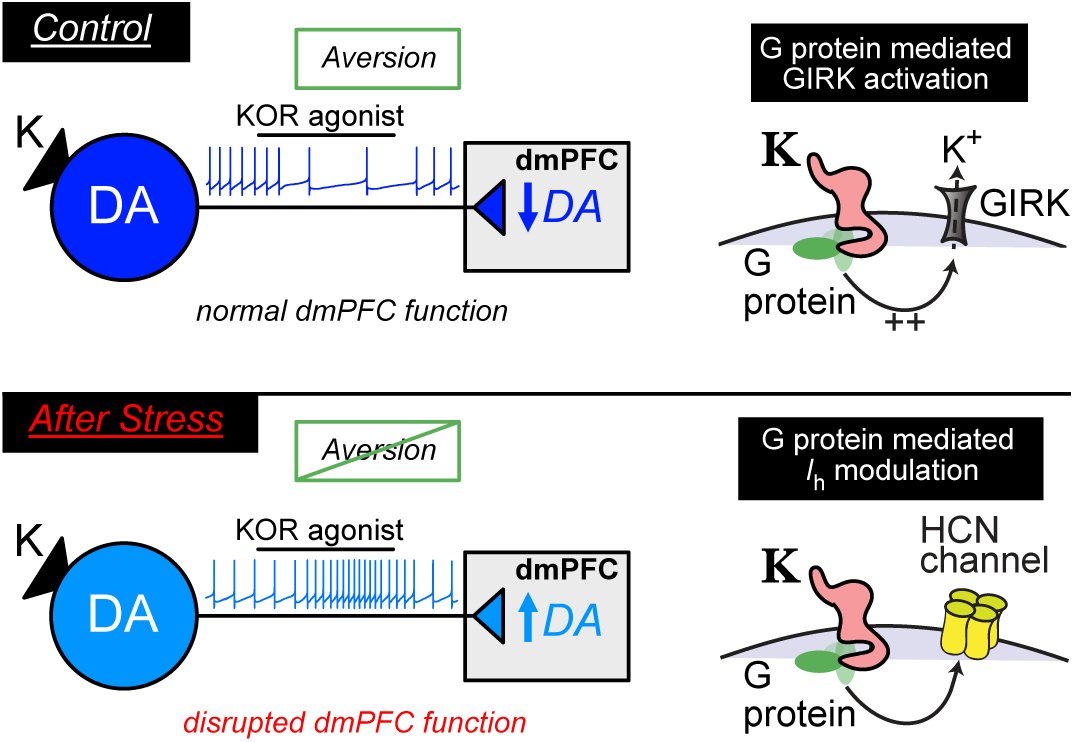
Summary: Aversive stress switches KOR signaling from GIRK mediated inhibitions to HCN channel mediated depolarizations. This switch specifically occurs in dmPFC-projecting VTA dopamine neurons.

The contributions of VTA dopamine neurons to the processing of aversive experiences encompass motivational and learning components and are circuit-specific (34). *In vivo* electrophysiology studies show that while many midbrain putative and opto-tagged dopamine neurons decrease their firing when a predicted reward is omitted (i.e., an aversive outcome) (35,36), some putative and directly identified dopamine neurons increase their firing in response to an aversive sensory stimulus (37,38). We previously demonstrated that in control rats, KOR activation inhibits VTA dopamine neurons that project to the mPFC and BLA, but not the NAc (18,19). With microdialysis, we confirmed that *in vivo* VTA KOR activation decreases dopamine release in mPFC but not in the NAc (19). Here we show a switch from inhibitory to excitatory signaling in a subpopulation of dmPFC-projecting VTA neurons after CRF exposure. This excitatory KOR effect may contribute to the reported stress induced activation of dmPFC-projecting VTA dopamine neurons (39,40). The loss of intra-VTA KOR agonist induced CPA after the stress induced switch in KOR signaling also implicates the dopaminergic projection to dmPFC in value signaling. Consistent with this, MOR activation in the VTA causes an increase in dopamine release in the anterior cingulate cortex (ACC, included in the dmPFC in this study), and lesioning dopamine terminals there decreases the magnitude of intra-VTA MOR agonist induced conditioned place preference (15). Also, genetic deletion of KOR from TH neurons and stress induced augmentation of cocaine CPP (41). The transfer function of dopamine activity in the mPFC has been extensively studied, but a synthesized understanding of its contributions to mPFC processing of value remains somewhat elusive (42–44).

Here we discovered a convergence of two neuropeptide systems, with CRF changing the sign of the physiological response of KORs in a specific neural circuit. There are at least a dozen possible sources of CRF to the VTA including the bed nucleus of the stria terminalis, dorsal raphe, parabrachial nucleus, lateral dorsal tegmentum, and local VTA neurons (45,46) any of which might be activated by aversive stressors. A subset of VTA neurons express CRF receptor 1 (CRF-R1) and the CRF binding protein, but not CRF receptor 2 (47,48). Blocking CRF-R1 in the VTA decreases foot shock stress induced reinstatement of cocaine seeking (39). Intra-VTA KOR blockade (13) and intra-dmPFC dopamine antagonists also mitigate stress induced reinstatement of cocaine seeking (39,49), a provocative parallel to the neuromodulatory circuit identified here.

We found that blocking *I*_h_ prevented KOR agonist induced depolarizations. Even though *I*_h_ is generated by hyperpolarization activated, cyclic nucleotide-gated (HCN) channels (50,51) and thus opioid receptors might interact with them though cAMP modulation, in the NRM KOR activation in tissue from control rats augments *I*_h_ through a cAMP-independent signaling pathway that involves intracellular release of Ca^2+^ (32). In these neurons, cAMP activation causes a positive shift in the voltage dependence of *I*_h_, but does not change the maximum *I*_h_ amplitude. In the VTA, the adenylyl cyclase activator forskolin causes a shift in the *I*_h_ V_1/2_ to a more depolarized potential (52). The isoform HCN3, which is expressed in the vast majority of VTA dopamine neurons and in many more VTA neurons than isoforms 1, 2 or 4 (52), is relatively insensitive to cAMP (50,53). Here, KOR activation after CRF did not alter *I*_h_ V_1/2_, consistent with a cAMP-independent effect.

An increase in HCN channel function in the VTA has been implicated in behavioral changes induced by aversive stressors. For instance, after social defeat stress in mice, *I*_h_ magnitude is larger in VTA neurons of susceptible mice compared to those of control mice (54). Blocking *I*_h_ in the VTA (54,55) and systemically (56) reverse the social avoidance induced by social defeat.

Our data suggest that *in vivo* activity at KORs in the VTA after stress increases *I*_h_ function in dmPFC-projecting dopamine neurons. Thus decreasing *I*_h_ function in the VTA may be one common therapeutic target, be it through direct HCN blockade or KOR antagonism.

What are the *in vivo* consequences of this switch in KOR signaling? At the circuit level, this switch means that the change in VTA dmPFC-projecting dopamine neuron firing will have an opposite sign when an endogenous KOR ligand is released into the VTA. That is, the peptidergic inputs to the VTA may have the same behavior-related firing patterns as before an aversive stressor, but the impact on dmPFC dopamine release will be the inverse. In addition to contributing to value encoding, dopamine function in the dmPFC also contributes to executive function (16,17). For instance, dopamine lesions in in the ACC impairs attentional set shifting (57) and dopamine receptor antagonists delivered to the prelimbic cortex degrade decision making (58). These and other studies suggest that ongoing or increases in dopamine release in the dmPFC contribute to saliency processing, and support working memory (59) which are particularly advantageous to engage following an aversive stressor. Following chronic stress that induces anhedonia, activating mPFC-projecting VTA dopamine neurons reverses immobility and loss of sucrose preference (60) and an increase in dopamine signaling in dmPFC also relieves ongoing pain (61). Provocatively, stress induced analgesia is also blocked by systemic administration of the KOR selective antagonist norBNI (2), which in the context of our results would prevent an increase in dopamine release in the dmPFC, although it is not yet known if dmPFC-projecting VTA dopamine neurons contribute to this phenomenon. Together, these studies raise the possibility that this switch in VTA KOR function is adaptive, with an optimal concentration of dopamine in the mPFC facilitating learning. Learning about acutely aversive experiences, and how to avoid them, is critical to survival.

## Acknowledgements

I thank Joseph R. Driscoll, Thomas J. Cirino, Peter Fong, Aashka K. Popat, Ben J. Ngu, Benjamin Snyder, Kasra Mansorian, and Madelyn Moulton for technical assistance. This work was supported by National Institutes of Health grants R01DA030529, R01DA042025 and R01DA008863, and funds from the State of California for medical research on alcohol and substance use. Manuscript was previously posted on BioRxiv (https://doi.org/10.1101/2025.08.09.669424). Data are available from OSF (doi:10.17605/OSF.IO/8FRHJ)

## Author Contributions

E.B.M. conceived of the research, designed the study, performed all electrophysiology and data analysis, generated the figures, and composed the manuscript.

## Disclosures

E.B.M. received consulting fees from Neumora Therapeutics, Inc.

## Supplementary Figure Captions

**Supplementary Figure 1.**
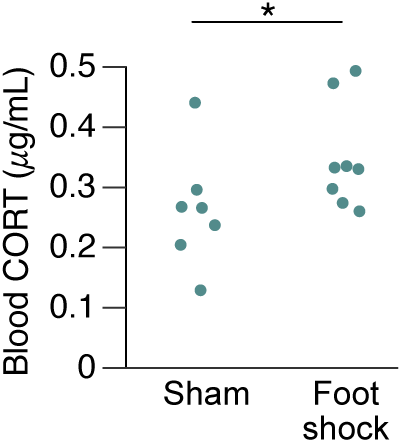
CORT measured (ELISA) from trunk blood samples collected from same day foot shock and sham animals during brain slice preparation. * *p* = 0.05

**Supplementary Figure 2.**
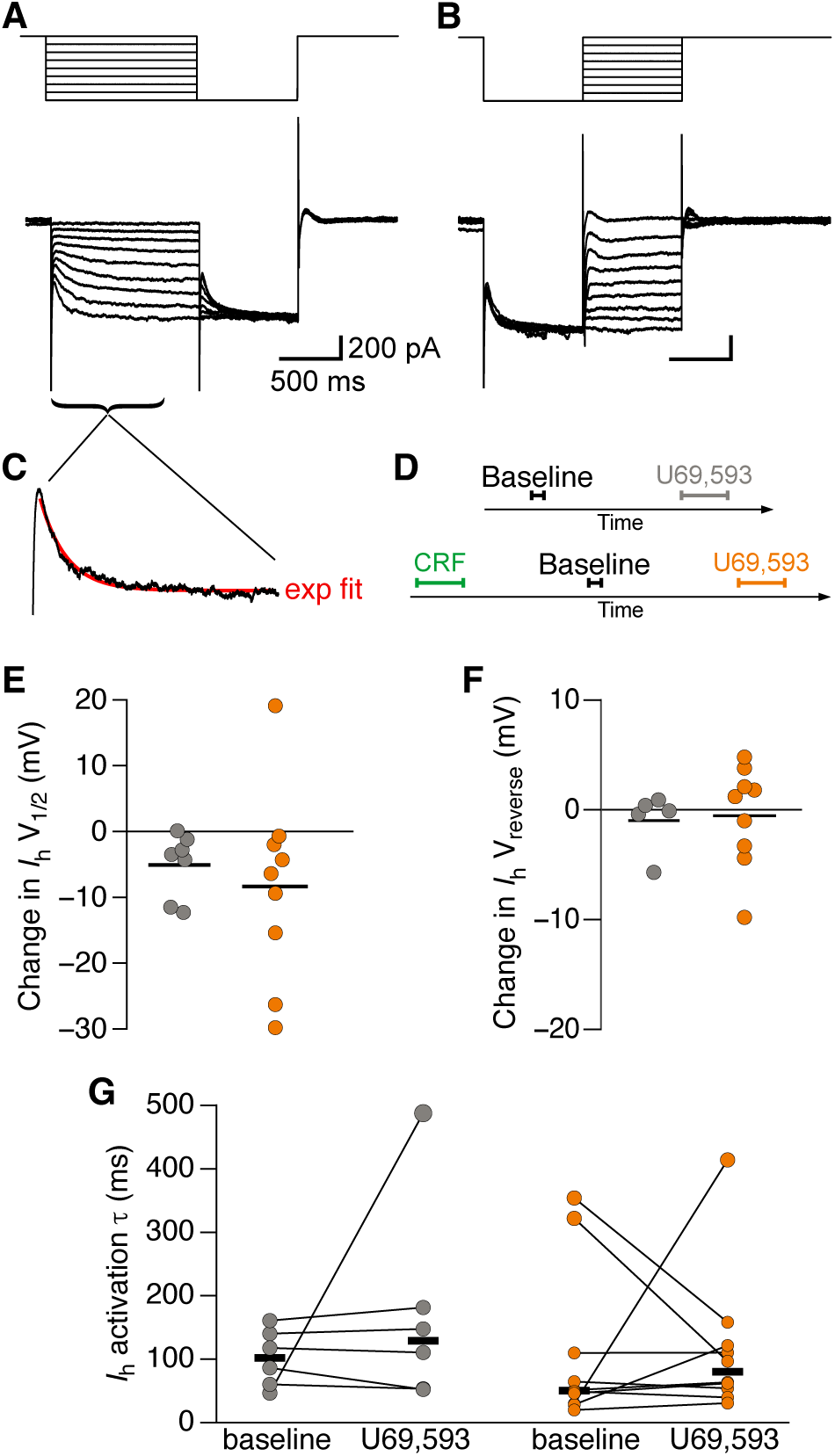
I_h_ measurements. (A) top, schematic of V step commands from V=-40 mV to quantify V_1/2_ and activation τ. Bottom, example response. (B) top, schematic of V step commands from V = −40 mV to quantify reversal V. Bottom, example response from the same neuron as in (A). (C) Example single exponential fit to data in (A). (D) Schematic of experiment design. Step series shown in (A) and (B) were applied to each neuron for a “baseline” measurement and again during U69,593 application. (E) Difference between V_1/2_ measured at baseline and in U69,593. Grey is no pretreatment, Orange is CRF pretreatment. (F) Difference between V_reverse_ measured at baseline and in U69,593. Grey is no pretreatment, Orange is CRF pretreatment. (G) *I*_h_ activation ι− at baseline vs in U69,593. Grey is no pretreatment, Orange is CRF pretreatment.

**Supplementary Figure 3.**
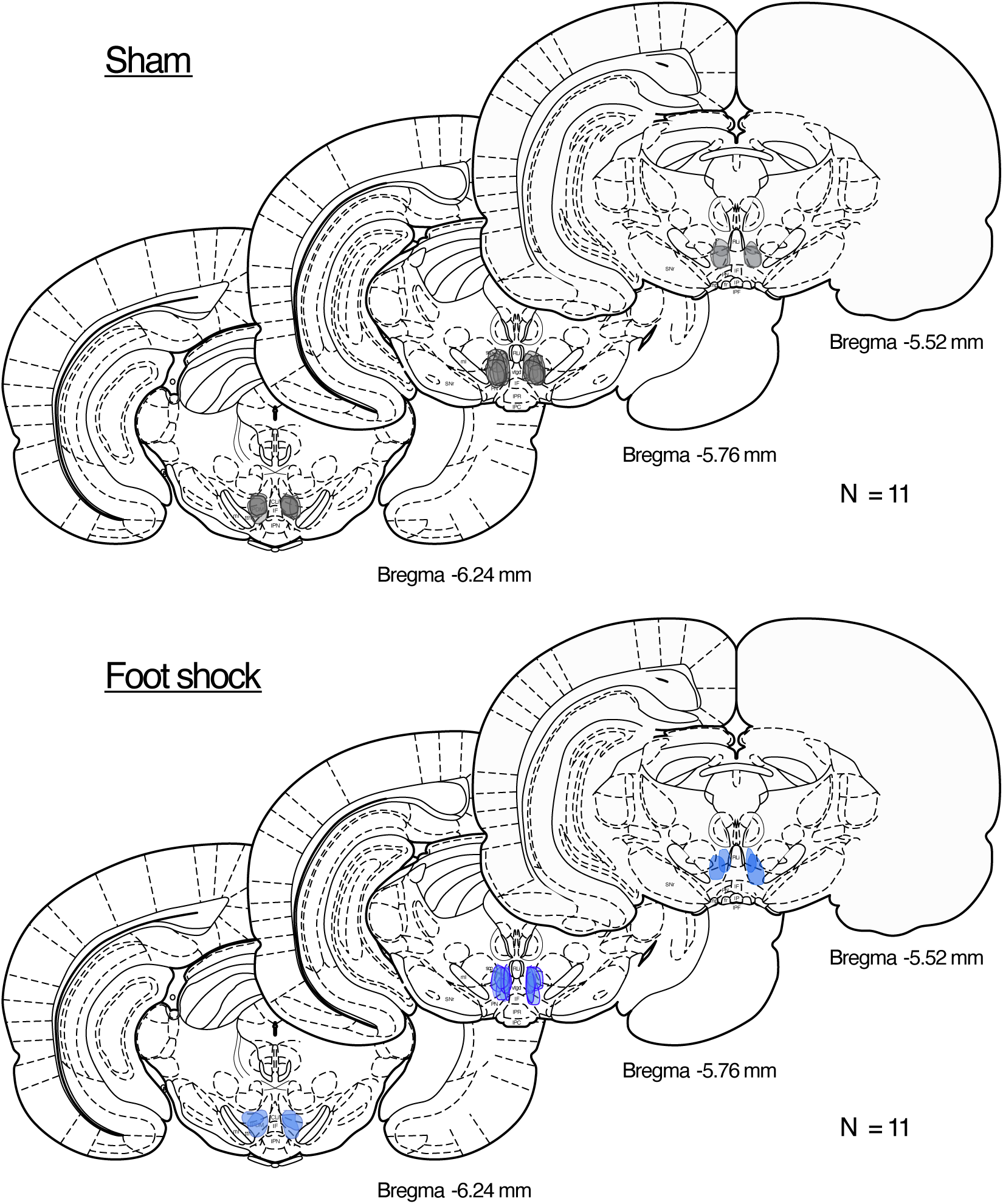
Microinjection sites for intra-VTA place conditioning experiments in Figure 5. Modified from (62).

**Supplementary Figure 4.**
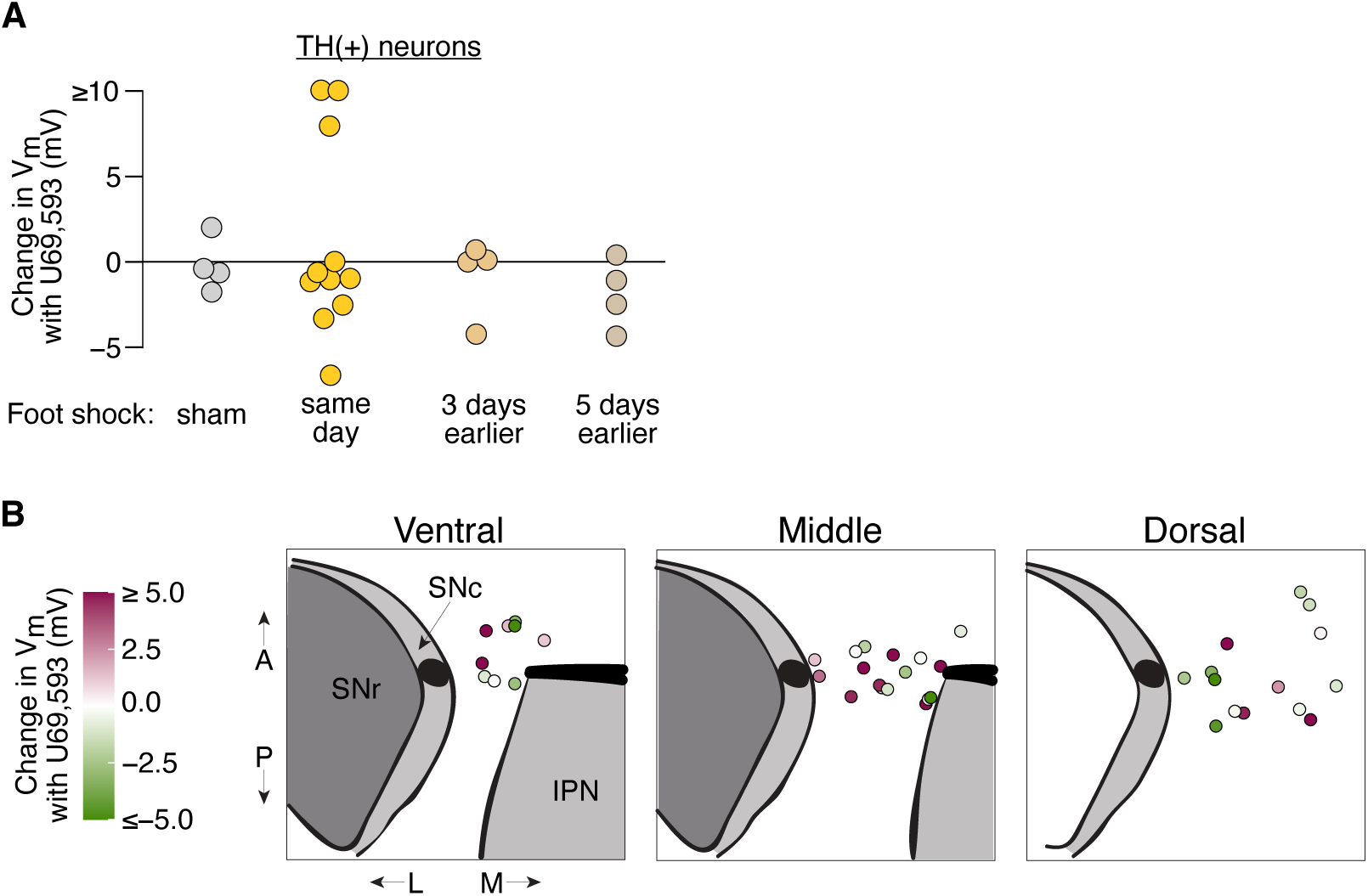
(A) Summary of responses to U69,593 in neurons identified at TH(+) with immunocytochemistry. These are a subset of the data in Fig. 1D. (B) Locations of VTA neurons recorded from same day foot shock stress rats. Color indicates change in Vm in response to U69,593.

**Supplementary Figure 5.**
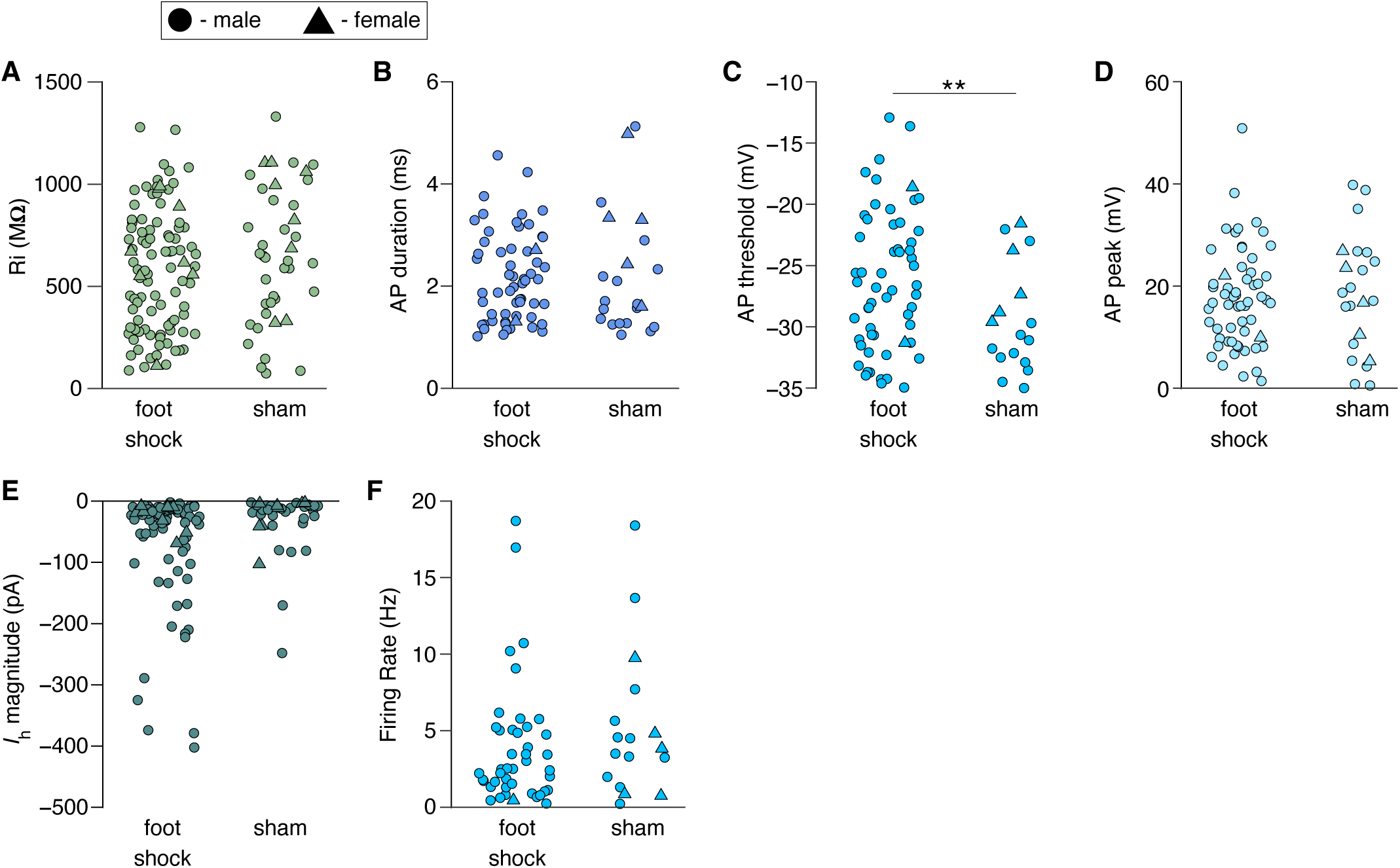
Physiological properties recorded in VTA neurons from foot shock (same day) or sham rats. Quantifications of data collected during the first 150 s immediately after whole cell access is achieved. (A) Ri, (B) AP duration, (C) AP threshold, (D) AP peak, (E) *I*_h_ magnitude (F) spontaneous firing rate. \*\**p* < 0.01

**Supplementary Figure 6.**
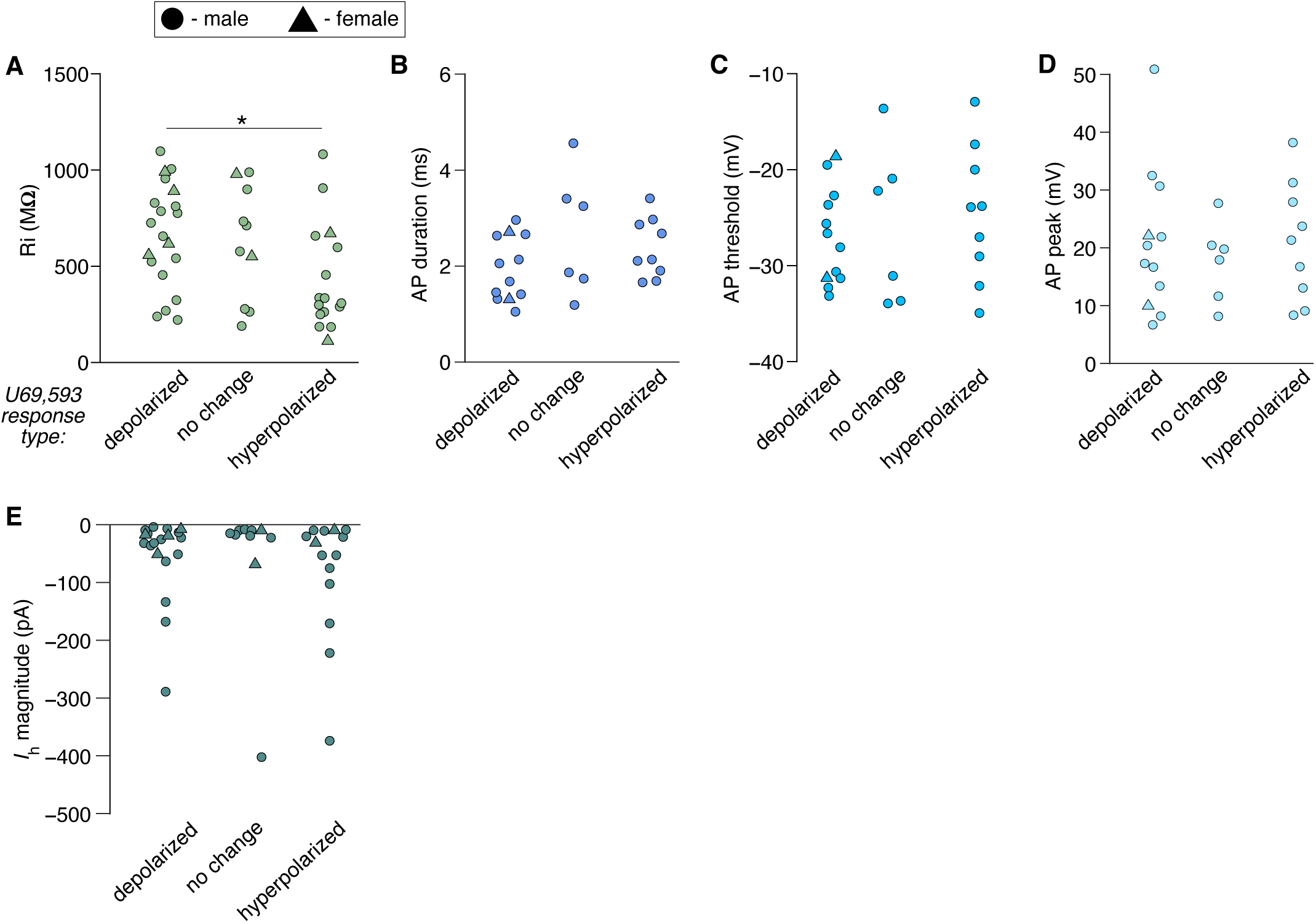
Foot shock rat data from Supplementary Figure 2, sorted by the type of response to U69,593. (A) Ri, (B) AP duration, (C) AP threshold, (D) AP peak, (E) *I*_h_ magnitude \**p* < 0.05

## Notes

### Summary of Updates

Corrected errors in the Reference section. Updated disclosures.

## References

1. Varlinskaya EI, Spear LP, Diaz MR (2018): Stress alters social behavior and sensitivity to pharmacological activation of kappa opioid receptors in an age-specific manner in Sprague Dawley rats. Neurobiol Stress 9: 124–132.

2. McLaughlin JP, Li S, Valdez J, Chavkin TA, Chavkin C (2006): Social defeat stress-induced behavioral responses are mediated by the endogenous kappa opioid system. Neuropsychopharmacology 31: 1241–8.

3. Massaly N, Copits BA, Wilson-Poe AR, Hipólito L, Markovic T, Yoon HJ, et al. (2019): Pain-Induced Negative Affect Is Mediated via Recruitment of The Nucleus Accumbens Kappa Opioid System. Neuron. 10.1016/j.neuron.2019.02.029

4. Groblewski PA, Zietz C, Willuhn I, Phillips PEM, Chavkin C (2015): Repeated stress exposure causes strain-dependent shifts in the behavioral economics of cocaine in rats. Addict Biol 20: 297–301.

5. Carlezon Jr. WA, Krystal AD (2016): Kappa-Opioid Antagonists for Psychiatric Disorders: From Bench to Clinical Trials. Depress Anxiety 33: 895–906.

6. Reeves KC, Shah N, Muñoz B, Atwood BK (2022): Opioid Receptor-Mediated Regulation of Neurotransmission in the Brain. Front Mol Neurosci 15: 919773.

7. Ji M, Gao Z, Yang J, Cai J, Li K, Wang J, et al. (2022): Dynorphin promotes stress-induced depressive behaviors by inhibiting ventral pallidal neurons in rats. Acta Physiol 236. 10.1111/apha.13882

8. Lemos JC, Roth CA, Messinger DI, Gill HK, Phillips PEM, Chavkin C (2012): Repeated stress dysregulates κ-opioid receptor signaling in the dorsal raphe through a p38α MAPK-dependent mechanism. J Neurosci Off J Soc Neurosci 32: 12325–12336.

9. Pan ZZ, Tershner SA, Fields HL (1997): Cellular mechanism for anti-analgesic action of agonists of the kappa-opioid receptor. Nature 389: 382–385.

10. Margolis EB, Hjelmstad GO, Bonci A, Fields HL (2003): Kappa-opioid agonists directly inhibit midbrain dopaminergic neurons. J Neurosci 23: 9981–6.

11. Bals-Kubik R, Ableitner A, Herz A, Shippenberg TS (1993): Neuroanatomical sites mediating the motivational effects of opioids as mapped by the conditioned place preference paradigm in rats. J Pharmacol Exp Ther 264: 489–95.

12. Ehrich JM, Messinger DI, Knakal CR, Kuhar JR, Schattauer SS, Bruchas MR, et al. (2015): Kappa Opioid Receptor-Induced Aversion Requires p38 MAPK Activation in VTA Dopamine Neurons. J Neurosci 35: 12917–31.

13. Graziane NM, Polter AM, Briand LA, Pierce RC, Kauer JA (2013): Kappa opioid receptors regulate stress-induced cocaine seeking and synaptic plasticity. Neuron 77: 942– 954.

14. Vijayraghavan S, Major AJ, Everling S (2017): Neuromodulation of Prefrontal Cortex in Non-Human Primates by Dopaminergic Receptors during Rule-Guided Flexible Behavior and Cognitive Control. Front Neural Circuits 11: 91.

15. Narita M, Matsushima Y, Niikura K, Narita M, Takagi S, Nakahara K, et al. (2010): Implication of dopaminergic projection from the ventral tegmental area to the anterior cingulate cortex in μ-opioid-induced place preference. Addict Biol 15: 434– 447.

16. Tichelaar JG, Sayalı C, Helmich RC, Cools R (2023): Impulse control disorder in Parkinson’s disease is associated with abnormal frontal value signalling. Brain J Neurol 146: 3676–3689.

17. Cools R, Froböse M, Aarts E, Hofmans L (2019): Dopamine and the motivation of cognitive control. Handb Clin Neurol 163: 123–143.

18. Margolis EB, Mitchell JM, Ishikawa J, Hjelmstad GO, Fields HL (2008): Midbrain dopamine neurons: projection target determines action potential duration and dopamine D(2) receptor inhibition. J Neurosci 28: 8908–13.

19. Margolis EB, Lock H, Chefer VI, Shippenberg TS, Hjelmstad GO, Fields HL (2006): Kappa opioids selectively control dopaminergic neurons projecting to the prefrontal cortex. Proc Natl Acad Sci U A 103: 2938–42.

20. Seutin V, Johnson SW, North RA (1994): Effect of dopamine and baclofen on N-methyl-D-aspartate-induced burst firing in rat ventral tegmental neurons. Neuroscience 58: 201–206.

21. Land BB, Bruchas MR, Lemos JC, Xu M, Melief EJ, Chavkin C (2008): The dysphoric component of stress is encoded by activation of the dynorphin kappa-opioid system. J Neurosci 28: 407–14.

22. Bruchas MR, Land BB, Lemos JC, Chavkin C (2009): CRF1-R activation of the dynorphin/kappa opioid system in the mouse basolateral amygdala mediates anxiety-like behavior. PloS One 4: e8528.

23. Van’t Veer A, Yano JM, Carroll FI, Cohen BM, Carlezon WA (2012): Corticotropin-releasing factor (CRF)-induced disruption of attention in rats is blocked by the κ-opioid receptor antagonist JDTic. Neuropsychopharmacol Off Publ Am Coll Neuropsychopharmacol 37: 2809–2816.

24. Wang B, Shaham Y, Zitzman D, Azari S, Wise RA, You ZB (2005): Cocaine experience establishes control of midbrain glutamate and dopamine by corticotropin-releasing factor: a role in stress-induced relapse to drug seeking. J Neurosci 25: 5389–96.

25. Xia Y-F, Margolis EB, Hjelmstad GO (2010): Substance P inhibits GABAB receptor signalling in the ventral tegmental area. J Physiol 588: 1541–1549.

26. Margolis EB, Hjelmstad GO, Fujita W, Fields HL (2014): Direct bidirectional μ-opioid control of midbrain dopamine neurons. J Neurosci 34: 14707–16.

27. Margolis EB, Fujita W, Devi LA, Fields HL (2017): Two delta opioid receptor subtypes are functional in single ventral tegmental area neurons, and can interact with the mu opioid receptor. Neuropharmacology 123: 420–432.

28. Cobb-Lewis DE, Sansalone L, Khaliq ZM (2023): Contributions of the Sodium Leak Channel NALCN to Pacemaking of Medial Ventral Tegmental Area and Substantia Nigra Dopaminergic Neurons. J Neurosci Off J Soc Neurosci 43: 6841–6853.

29. Khaliq ZM, Bean BP (2010): Pacemaking in dopaminergic ventral tegmental area neurons: depolarizing drive from background and voltage-dependent sodium conductances. J Neurosci Off J Soc Neurosci 30: 7401–7413.

30. Philippart F, Khaliq ZM (2018): Gi/o protein-coupled receptors in dopamine neurons inhibit the sodium leak channel NALCN. eLife 7: e40984.

31. Ingram SL, Williams JT (1994): Opioid inhibition of Ih via adenylyl cyclase. Neuron 13: 179–186.

32. Pan ZZ (2003): κ-Opioid receptor-mediated enhancement of the hyperpolarization-activated current (Ih) through mobilization of intracellular calcium in rat nucleus raphe magnus. J Physiol 548: 765–775.

33. Bruchas MR, Land BB, Aita M, Xu M, Barot SK, Li S, Chavkin C (2007): Stress-induced p38 mitogen-activated protein kinase activation mediates kappa-opioid-dependent dysphoria. J Neurosci 27: 11614–23.

34. Morales M, Margolis EB (2017): Ventral tegmental area: cellular heterogeneity, connectivity and behaviour. Nat Rev Neurosci. 10.1038/nrn.2016.165

35. Cohen JY, Haesler S, Vong L, Lowell BB, Uchida N (2012): Neuron-type-specific signals for reward and punishment in the ventral tegmental area. Nature 482: 85–8.

36. Schultz W (1998): Predictive reward signal of dopamine neurons. J Neurophysiol 80: 1– 27.

37. Brischoux F, Chakraborty S, Brierley DI, Ungless MA (2009): Phasic excitation of dopamine neurons in ventral VTA by noxious stimuli. Proc Natl Acad Sci U A 106: 4894–9.

38. Bromberg-Martin ES, Matsumoto M, Hikosaka O (2010): Dopamine in motivational control: rewarding, aversive, and alerting. Neuron 68: 815–34.

39. Vranjkovic O, Van Newenhizen EC, Nordness ME, Blacktop JM, Urbanik LA, Mathy JC, et al. (2018): Enhanced CRFR1-Dependent Regulation of a Ventral Tegmental Area to Prelimbic Cortex Projection Establishes Susceptibility to Stress-Induced Cocaine Seeking. J Neurosci Off J Soc Neurosci 38: 10657–10671.

40. Deutch AY, Lee MC, Gillham MH, Cameron DA, Goldstein M, Iadarola MJ (1991): Stress selectively increases fos protein in dopamine neurons innervating the prefrontal cortex. Cereb Cortex N Y N 1991 1: 273–292.

41. Abraham AD, Casello SM, Land BB, Chavkin C (2022): Optogenetic stimulation of dynorphinergic neurons within the dorsal raphe activate kappa opioid receptors in the ventral tegmental area and ablation of dorsal raphe prodynorphin or kappa receptors in dopamine neurons blocks stress potentiation of cocaine reward. Addict Neurosci 1: 100005.

42. Sheynikhovich D, Otani S, Bai J, Arleo A (2023): Long-term memory, synaptic plasticity and dopamine in rodent medial prefrontal cortex: Role in executive functions. Front Behav Neurosci 16: 1068271.

43. Durstewitz D, Seamans JK (2008): The dual-state theory of prefrontal cortex dopamine function with relevance to catechol-o-methyltransferase genotypes and schizophrenia. Biol Psychiatry 64: 739–749.

44. Robinson SE, Sohal VS (2017): Dopamine D2 Receptors Modulate Pyramidal Neurons in Mouse Medial Prefrontal Cortex through a Stimulatory G-Protein Pathway. J Neurosci Off J Soc Neurosci 37: 10063–10073.

45. Soden ME, Yee JX, Cuevas B, Rastani A, Elum J, Zweifel LS (2022): Distinct Encoding of Reward and Aversion by Peptidergic BNST Inputs to the VTA. Front Neural Circuits 16: 918839.

46. Grieder TE, Herman MA, Contet C, Tan LA, Vargas-Perez H, Cohen A, et al. (2014): VTA CRF neurons mediate the aversive effects of nicotine withdrawal and promote intake escalation. Nat Neurosci 17: 1751–1758.

47. Van Pett K, Viau V, Bittencourt JC, Chan RKW, Li H-Y, Arias C, et al. (2000): Distribution of mRNAs encoding CRF receptors in brain and pituitary of rat and mouse. J Comp Neurol 428: 191–212.

48. Wang H-L, Morales M (2008): Corticotropin-releasing factor binding protein within the ventral tegmental area is expressed in a subset of dopaminergic neurons. J Comp Neurol 509: 302–318.

49. McFarland K, Kalivas PW (2001): The circuitry mediating cocaine-induced reinstatement of drug-seeking behavior [no. 21]. J Neurosci Off J Soc Neurosci 21: 8655–8663.

50. Combe CL, Gasparini S (2021): Ih from synapses to networks: HCN channel functions and modulation in neurons. Prog Biophys Mol Biol 166: 119–132.

51. Monteggia LM, Eisch AJ, Tang MD, Kaczmarek LK, Nestler EJ (2000): Cloning and localization of the hyperpolarization-activated cyclic nucleotide-gated channel family in rat brain. Mol Brain Res 81: 129–139.

52. Mu L, Liu X, Yu H, Vickstrom CR, Friedman V, Kelly TJ, et al. (2023): cAMP-mediated upregulation of HCN channels in VTA dopamine neurons promotes cocaine reinforcement. Mol Psychiatry 28: 3930–3942.

53. Mistrík P, Mader R, Michalakis S, Weidinger M, Pfeifer A, Biel M (2005): The Murine HCN3 Gene Encodes a Hyperpolarization-activated Cation Channel with Slow Kinetics and Unique Response to Cyclic Nucleotides. J Biol Chem 280: 27056– 27061.

54. Cao J-L, Covington HE, Friedman AK, Wilkinson MB, Walsh JJ, Cooper DC, et al. (2010): Mesolimbic dopamine neurons in the brain reward circuit mediate susceptibility to social defeat and antidepressant action. J Neurosci Off J Soc Neurosci 30: 16453– 16458.

55. Cai M, Zhu Y, Shanley MR, Morel C, Ku SM, Zhang H, et al. (2023): HCN channel inhibitor induces ketamine-like rapid and sustained antidepressant effects in chronic social defeat stress model. Neurobiol Stress 26: 100565.

56. Teichman EM, Hu J, Lin H, Fisher-Foye RL, Blando A, Hu X, et al. (2025): Design and validation of novel brain-penetrant HCN channel inhibitors to ameliorate social stress-induced susceptible phenotype. Mol Psychiatry. 10.1038/s41380-025-02972-8

57. Mk C, Cs P, Ja M (2024): Dopaminergic lesions of the anterior cingulate cortex of rats increase vulnerability to salient distractors. Eur J Neurosci 59. 10.1111/ejn.16352

58. Jenni NL, Larkin JD, Floresco SB (2017): Prefrontal Dopamine D1 and D2 Receptors Regulate Dissociable Aspects of Decision Making via Distinct Ventral Striatal and Amygdalar Circuits. J Neurosci 37: 6200–6213.

59. Ott T, Nieder A (2019): Dopamine and Cognitive Control in Prefrontal Cortex. Trends Cogn Sci 23: 213–234.

60. Liu D, Tang Q-Q, Wang D, Song S-P, Yang X-N, Hu S-W, et al. (2020): Mesocortical BDNF signaling mediates antidepressive-like effects of lithium. Neuropsychopharmacology 45: 1557–1566.

61. K L, C Q, E N, F P, P S (2021): Decreased dopaminergic inhibition of pyramidal neurons in anterior cingulate cortex maintains chronic neuropathic pain. Cell Rep 37. 10.1016/j.celrep.2021.109933

62. Paxinos G, Watson C (1997): The Rat Brain in Stereotaxic Coordinates, Compact, 3rd ed. San Diego: Academic Press.

